# The Memorability of Voices is Predictable and Consistent across Listeners

**DOI:** 10.1101/2024.02.08.579540

**Authors:** Cambria Revsine, Esther Goldberg, Wilma A. Bainbridge

**Author notes:** Correspondence to: Cambria Revsine, 301 Green Hall, 5848 S. University Ave., Chicago, IL 60637.

## Abstract

Memorability, the likelihood that a stimulus is remembered, is an intrinsic stimulus property that is highly consistent across people—participants tend to remember and forget the same faces, objects, and more. However, these consistencies in memory have thus far only been observed for visual stimuli. We provide the first study of auditory memorability, collecting recognition memory scores from over 3000 participants listening to a sequence of different speakers saying the same sentence. We found significant consistency across participants in their memory for voice clips and for speakers across different utterances. Next, we tested regression models incorporating both low-level (e.g., fundamental frequency) and high-level (e.g., dialect) voice properties to predict their memorability. These models were significantly predictive, and cross-validated out-of-sample, supporting an inherent memorability of speakers’ voices. These results provide the first evidence that listeners are similar in the voices they remember, which can be reliably predicted by quantifiable voice features.

## Introduction

We can all think of certain voices that stick in our minds long after we hear them— perhaps the voice of a famous celebrity, or the voice of a stranger with whom we strike up a brief conversation. For example, most people would agree that Morgan Freeman’s voice is easier to recall than Brad Pitt’s, despite both being incredibly famous actors. But what makes certain speakers more memorable than others? And is this largely dependent on an individual’s past experiences, or do the same types of voices tend to be remembered and forgotten across people?

Although the majority of memory research has been carried out in the visual domain, this area of audition is equally deserving of study. Our everyday experiences do not just contain visual input, but are richly multisensory and dynamic. Auditory information, particularly the human voice, is thus integral to many cognitive processes, including communication, learning, and memory.

A *memorability* approach, which bridges the fields of perception and memory, can help address these open questions. Memorability is an intrinsic property of stimuli that quantifies how likely a given stimulus is to be remembered (Bainbridge et al., 2013; Isola et al., 2011). Critically, this property is highly consistent across the population—in other words, people tend to remember and forget the same images as one another. Memorability effects have been identified in many visual stimulus types, including faces (Bainbridge et al., 2013), scenes (Isola et al., 2011, 2014), objects (Kramer et al., 2023), words (Xie et al., 2020), infographics (Borkin et al., 2013), artwork (Davis & Bainbridge, 2023), and even dance moves (Ongchoco et al., 2023). In addition to static images, the memorability of faces has been shown to be consistent across different expressions and viewpoints (Bainbridge, 2017). Regression models containing human-labeled image features have been able to explain, at most, about half of the variance in visual memorability (Bainbridge et al., 2013; Kramer et al., 2023). These studies and others have demonstrated a greater influence of semantic, or high-level, features (e.g., animacy) over perceptual, low-level features (e.g., color) in determining this image property (Kramer et al., 2023; Needell & Bainbridge, 2022; Xie et al., 2020).

Up to this point, however, memorability research has been carried out entirely with visual stimuli. Thus, it is still unknown whether these properties of visual memorability also apply in other sensory domains. In this study, we aimed to evaluate, for the first time, whether there are consistencies in the auditory stimuli people remember as well. We chose to investigate voices specifically, as they contain both low-level properties, like pitch, and high-level properties, like dialect, which may differentially affect their memorability, as is the case with visual stimuli. There is reason to believe that intrinsic memorability would exist for voices. Recognition accuracy for speakers is fairly high (Palmeri et al., 1993), suggesting that some speakers may be consistently better remembered than others. Like faces, voices are highly relevant social stimuli, and there are proposed parallels in the functional and neural organization of both stimulus types (Belin et al., 2004; Young et al., 2020). Furthermore, subjective personality judgments of both faces and voices have been shown to be consistent across observers (McAleer et al., 2014; Mileva & Lavan, 2023; Todorov et al., 2008; but see Tompkinson et al., 2023). Such traits significantly predict the memorability of faces (Bainbridge et al., 2013).

However, there is also considerable evidence suggesting that auditory memorability effects may be less consistent than visual memorability, or even nonexistent. Compared with humans’ robust long-term memory for naturalistic images (Brady et al., 2008; Standing, 1973), memory for natural sounds and speech is thought to be inferior (Bigelow & Poremba, 2014; Cohen et al., 2009; Fritz et al., 2005). Research on earwitness testimony echoes this finding, as performance on voice lineup tasks is generally poor, especially with limited exposure to the voices (Clifford, 1980; Yarmey et al., 1994). This behavioral difference may be explained by the fact that there is limited evidence for auditory sensitivity in memory-related brain regions like the perirhinal cortex (Bigelow & Poremba, 2014; Munoz-Lopez et al., 2010; Peters et al., 2007). Finally, there are fundamental differences between voices and faces which may differentially affect their memorability. Voices are inherently more dynamic and carry both identity and speech information, so additional processes are involved in recognizing speakers (Magnuson et al., 2021).

In order to test these two competing hypotheses, we ran three experiments in which over 3000 participants were tested on their recognition memory performance for hundreds of speakers’ utterances. We found the first evidence that the memorability of voice clips is consistent across listeners, and also that the memorability of speakers themselves is consistent across different spoken sentences. We also aimed to predict this memorability from a mix of objective, low-level features and subjective, high-level features, the latter ratings collected from a separate group of nearly 1000 participants. Regression models containing these features were able to significantly predict the memorability of the voice stimuli, and successfully cross-validated out-of-sample. These results reveal that, in addition to memory for images, peoples’ memory for dynamic auditory stimuli is highly consistent, and can be reliably predicted by a combination of acoustic voice features.

## Results

In this experiment, participants (N=3340, Amazon Mechanical Turk) performed a 10-minute continuous recognition task in which they heard a randomized sequence of different speakers speaking a single sentence, with many speakers repeated once throughout the task. Three versions of the experiment were run on separate groups of participants, which differed in the sentence used as stimuli and/or the task demands. Experiment 1 contained stimuli of one sentence (“*She had your dark suit in greasy wash water all year*”) and Experiment 2 contained stimuli of another sentence (“*Don’t ask me to carry an oily rag like that*”). In both of these experiments, participants indicated with a key press whenever they heard a repeated voice clip. Experiment 3 contained a mix of both sentences. In this experiment, “repeats” were always of the same speaker speaking the other sentence, and participants indicated whenever they heard a repeated *speaker*. Repeated stimuli could be fillers, which repeated after a few trials and were used as a vigilance check, or targets, which repeated on average 20 trials (140 seconds) after the first presentation in Experiments 1 and 2 (**Figure 1**). In Experiment 3, targets repeated approximately 12 trials (84 seconds) after the first presentation, to account for the increased difficulty of the task. Each target repeat was heard by an average of 43 participants in Exp. 1, 47 participants in Exp. 2, and 48 participants in Exp. 3.

**Figure 1.**
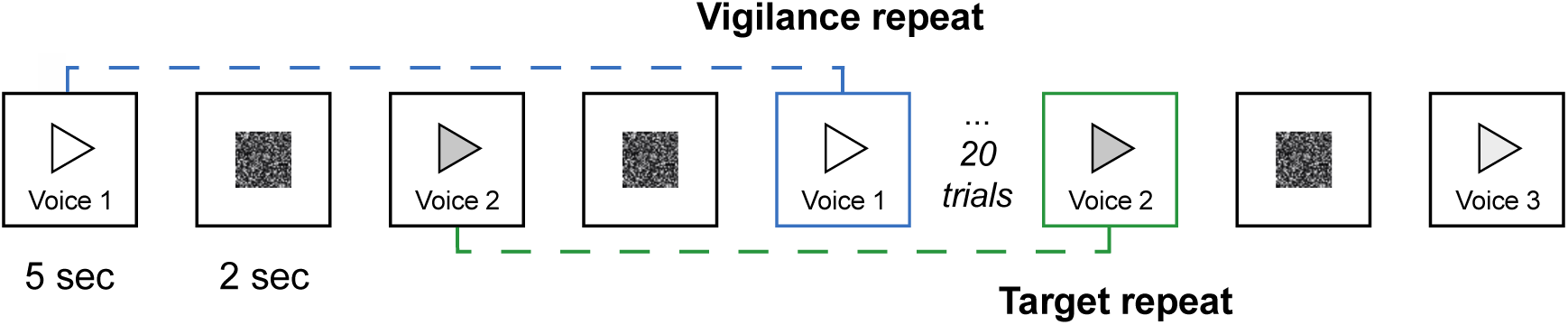
Memory experiment paradigm. Participants heard a sequence of voice clips of different speakers speaking a single sentence (each clip lasting up to five seconds), and pressed a key whenever they heard a repeated speaker. In Experiments 1-2 (shown here), vigilance repeats occurred after approximately five trials (∼ 35 seconds) and target repeats occurred after approximately 20 trials (∼ 140 seconds). In Experiment 3, repeats occurred slightly closer together.

### Accuracy and consistency of memory performance for voice clips

We first aimed to characterize memory for voice clips when speakers repeated the same utterances, in Experiments 1 and 2. For all stimuli, we measured the hit rate (HR) and false alarm rate (FAR) from participants’ memory performance when the voice clip was a target stimulus. Average HR for Sentence 1 stimuli in Exp. 1 was 0.53 (SD = 0.1), and average HR for Sentence 2 stimuli in Exp. 2 was 0.55 (SD = 0.09). Average FAR for Sentence 1 was 0.42 (SD = 0.1), and average FAR for Sentence 2 was 0.42 (SD = 0.1). From these two measures, we calculated the *d’* of each voice clip, which served as our main metric of memorability. Average *d’* for Sentence 1 was 0.28 (SD = 0.3) and average *d’* for Sentence 2 was 0.34 (SD = 0.29).

To address our central question of whether the memorability of voices is consistent across listeners, we ran a split-half consistency analysis on *d’* scores of target voice clips over 1000 iterations (see Methods). Briefly, if correlations between memorability scores from random splits of the participant pool are significantly above chance, this suggests that participants are similar in their memory, and false memory, for stimuli. We observed significant consistency in *d’* for both Sentence 1 (Spearman-Brown corrected ρ = 0.29, *p* = 0.002) and Sentence 2 (Spearman-Brown corrected ρ = 0.27, *p* = 0.002) (**Figure 2a-b**). Thus, in two separate sets of stimuli, we found that the same voice clips tended to be remembered and forgotten across listeners.

**Figure 2.**
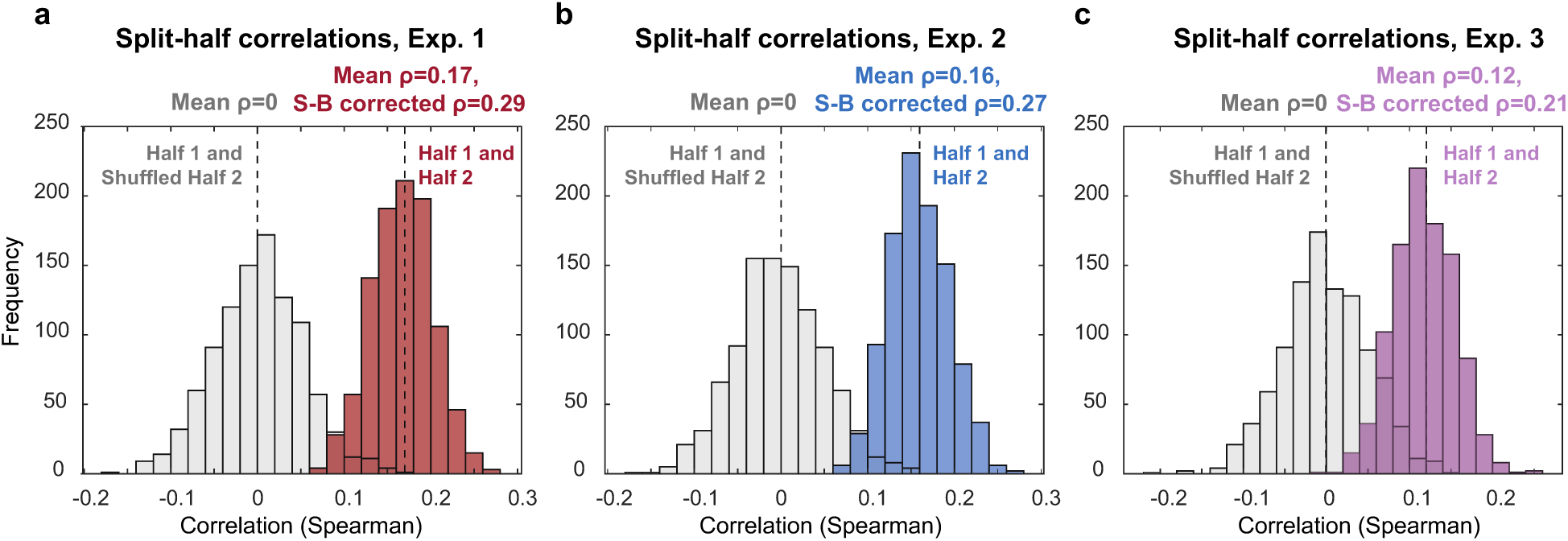
a) Split-half consistency analysis results for Sentence 1 voice clips in Experiment 1. The distribution of Spearman correlation coefficients between stimulus *d’* scores from random halves of the participant pool, over 1000 iterations, is shown in red. The chance-level distribution of correlations between a random half of participants and a shuffled other half, also over 1000 iterations, is shown in gray. Dashed lines indicate the average correlation value of each distribution. b) Split-half consistency analysis results for Sentence 2 voice clips in Experiment 2. Layout as in a. c) Split-half consistency analysis results for Experiment 3. Here, stimulus *d’* scores were calculated for repeats of the same speakers across different sentences. Layout as in a and b.

### Memory for speakers across utterances

Because the same speakers provided the stimuli in Experiment 1 and 2, we tested for consistency in the memorability of speakers across utterances. In other words, were the most memorable and forgettable speakers of Sentence 1 also the most memorable and forgettable speakers of Sentence 2? We observed a significant correlation between the *d’* scores of the Experiment 1 and 2 stimuli (ρ = 0.17, *p* < 0.001) (**Figure 3a**). This suggests that there is consistent memorability of the voices of speakers, regardless of what the speakers are saying.

**Figure 3.**
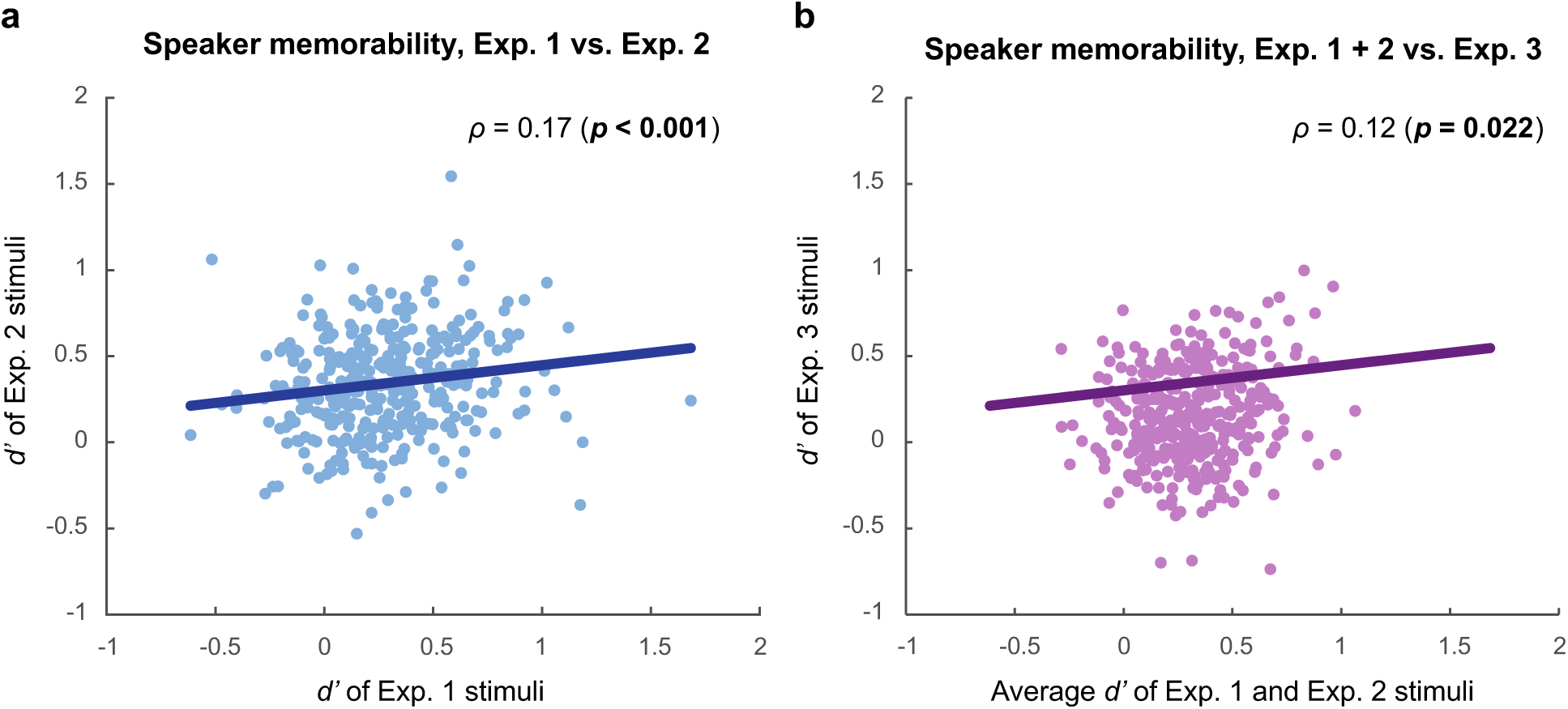
a) Correlation between memorability scores from Experiment 1 and Experiment 2. *d’* of Experiment 1 stimuli is plotted along the x-axis, and *d’* of Experiment 2 stimuli—the same speakers speaking the other sentence—is plotted along the y-axis. Line of best fit shown in dark blue. b) Comparison of memorability scores from all three experiments. Average *d’* of Experiment 1 and 2 stimuli is plotted along the x-axis, and *d’* of Experiment 3 stimuli—the same speakers speaking both sentences—is plotted along the y-axis. Line of best fit shown in dark purple.

These results provided evidence supporting the existence of speaker-level memorability. However, to more directly test this, we repeated the same analyses on the results of Experiment 3, in which participants identified different utterances from the same speakers. A split-half consistency analysis on *d’* scores of the target repeats in Experiment 3 also revealed a significant correlation (Spearman-Brown corrected ρ = 0.21, *p* = 0.012) (**Figure 2c**). Therefore, participants were consistent in their memory performance not just for individual voice clips, but for speakers themselves.

Lastly, we compared the memorability of speakers across all three experiments. We first computed the average *d’* of speakers across the sentences in Experiment 1 and 2, and then correlated them with the *d’* scores of the same speakers from Experiment 3 where they spoke both sentences. This resulted in a significant correlation of 0.12 (*p* = 0.022) (**Figure 3b**). This finding provides yet more evidence for the similarity of memory performance for speakers’ voices across separate groups of participants and different utterances.

### Characterizing low- and high-level voice features

Having established the robust memorability effect of our voice stimuli, we next wondered whether, and how well, we could predict this memorability from a range of both low- and high-level voice features. Using the audio software VoiceSauce (Shue et al., 2011), we measured the mean and standard deviation of 44 low-level acoustic features of each voice clip. These included measures such as the fundamental frequency (base pitch), frequency and bandwidth of formants, and amplitude of harmonics (see Methods for full list). We also included the duration of the voice clips (i.e., the speed of speech) in the low-level features category. For all 89 low-level measures, we observed a significant correlation between the measures of Sentence 1 and 2 stimuli of the same speakers (all *p* < 0.05).

In addition to these objective, low-level features, we also investigated the contribution of subjective, high-level features. We collected a list of 14 personality-related adjectives and their antonyms, such as *aggressive/calm* and *confident/uncertain* (see Methods for all attributes), borrowed from previous work on memorability and personality judgments of faces (Bainbridge et al., 2013; Oosterhof & Todorov, 2008; Vokey & Read, 1992) and personality judgments of voices (McAleer et al., 2014). The attributes and their antonyms were randomly split across two versions of a survey, taken by a new group of AMT participants (N=903). In the survey, participants rated Sentence 1 voice clips on a scale from 1-9 according to all attributes (e.g., “How *confident* does this voice sound?”). We first correlated ratings for each attribute-antonym pair. Surprisingly, only *masculine/feminine* (*r* = -0.12, *p* < 0.001) and *calm/aggressive* (*r* = -0.03, *p* = 0.02) were significantly negatively correlated. Suspecting that this might be due to inconsistency in the ratings, we aligned ratings of antonym pairs onto the same scale, and ran a split-half consistency analysis on all 30 ratings of each attribute. *Calm/aggressive* (Spearman-Brown corrected ρ = 0.20, *p* = 0.02), *expressive/unemotional* (Spearman-Brown corrected ρ = 0.17, *p* = 0.03), *masculine/feminine (*Spearman-Brown corrected ρ = 0.83, *p* < 0.001), and *young/old* (Spearman-Brown corrected ρ = 0.23, *p* = 0.003) were the only attributes to show significant consistency in participants’ ratings. Thus, it appears that participants generally did not agree on their ratings of these subjective, high-level properties for a speaker’s voice. For this reason, we decided not to collect ratings of the Sentence 2 stimuli.

Regardless, we then investigated the relationships between both the low-level and high-level features by correlating all pairs of features. As can be seen in the correlation matrix of the Sentence 1 measures in **Figure 4** (left), many low-level features were highly correlated with each other (*r* > 0.7 or < -0.7). This was expected as many features are closely related, e.g., the amplitude of harmonics (H1-H4, A1-A3), and some features are computed multiple times using different algorithms, e.g., the fundamental frequency (F0). Interestingly, correlations among high-level features (**Figure 4**, right) were generally not as strong as those between low-level features. However, many of these correlations are sensical; for example, we observed fairly high positive correlations (*r* > 0.6) between *confident* and *trustworthy*, *likable*, and *competent*, and fairly high negative correlations (*r* ∼= -0.4) between *warm* and *dominant*, and *young* and *masculine*. Only a single high-level feature (the *masculine/feminine* attribute) showed high correlations with (several) low-level features. Although this matrix only contains results of the Sentence 1 features, correlations between low-level features of the Sentence 2 stimuli were qualitatively similar (**Supplementary Figure 1**).

**Figure 4.**
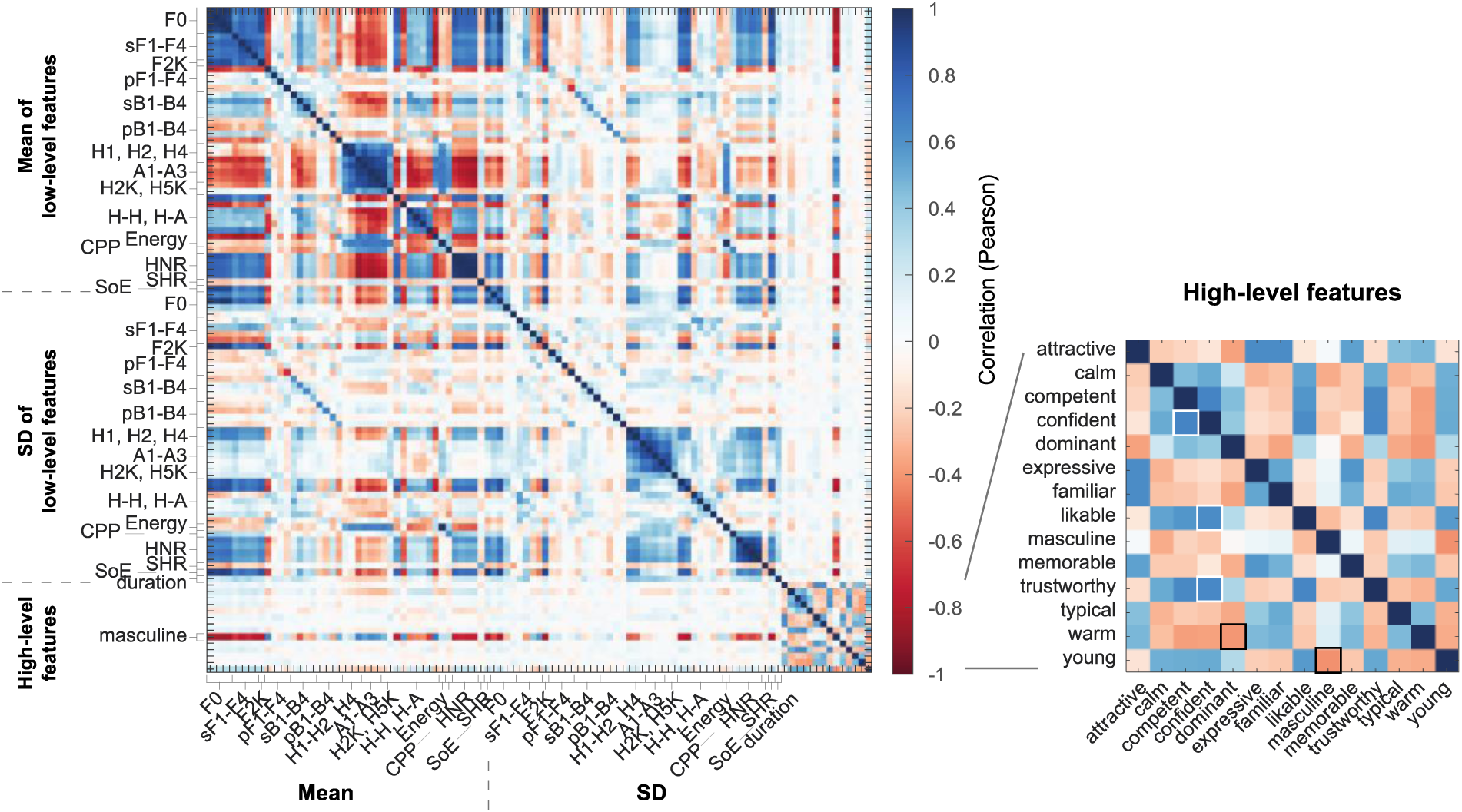
Correlation matrix of Sentence 1 voice features. As seen along the axes, the full matrix on the left contains mean measures of the 44 low-level acoustic features, followed by the standard deviation (SD) of the same features, followed by the “duration” measure, and finally the high-level personality-related attributes. Many strong correlations can be seen between related low-level features. Only one high-level feature (the *masculine* rating) was significantly correlated with low-level features. Correlations between high-level features (inset on right) were generally lower than those between low-level features; however, examples of relatively high correlations between certain attribute pairs are highlighted. Note: F0 includes strF0, sF0, pF0, and shrF0. H-H, H-A includes H1-H2, H2-H4, H1-A1, H1-A2, H1-A3, H4-H2K, and H2K-H5K. HNR includes HNR05, HNR15, HNR25, and HNR35.

### Predicting voice memorability from auditory features

After quantifying the low- and high-level features of the voice clips, we aimed to see whether we could significantly predict their memorability scores from a combination of these features, and if so, how much variance in memorability we could explain. We used stepwise linear regression to arrive at two multiple regression models, one predicting *d’* of the Sentence 1 stimuli in Experiment 1, and the other predicting *d’* of the Sentence 2 stimuli in Experiment 2. For the Sentence 1 model, we considered all 103 features described above (89 low-level and 14 high-level), as well as the gender and dialect of speakers. For the Sentence 2 model, we only considered the 89 low-level features, along with gender and dialect, since ratings of the subjective high-level features were not collected for these stimuli.

The final Sentence 1 model was significantly predictive of voice clip *d’* (*F*(7,370) = 13.8, *p* < 0.001), and had an adjusted *R^2^* value of 0.09 (**Supplementary Table 1**). The final Sentence 2 model was significantly predictive of *d’* as well (*F*(7,370) = 10.1, *p* < 0.001), with an adjusted *R^2^* value of 0.15 (**Supplementary Table 2**). Though these *R^2^* values may seem low, we can compare them to ceiling performance by dividing them by the Spearman-Brown corrected split-half correlations on the *d’* scores (**Figure 2**) to obtain a more accurate estimate of model performance. The Sentence 1 model therefore explained 31% of the variance ceiling, and the Sentence 2 model explained 56% of the variance ceiling. Both models contained only low-level, acoustic features as predictors (see **Supplementary Tables** for information on individual predictors).

We were interested in seeing whether these models could cross-validate to predict the memorability of the same speakers speaking the other sentence. To test this, we took the Sentence 1 model, which had been trained on the Sentence 1 voice features, and tested it on the Sentence 2 measures of the same features to predict Sentence 2 *d’*. We then correlated the predicted and true Sentence 2 *d’* scores. Doing so resulted in a significant correlation of 0.14 (*p* = 0.007). We then did the same using the Sentence 2 model to predict Sentence 1 *d’.* The predicted and true Sentence 1 memorability scores were also significantly correlated (*r* = 0.16, *p* = 0.001). Since both models successfully predicted the memorability of the other sentence, this suggests that there is intrinsic memorability of the voices of speakers, regardless of the spoken content.

Finally, we ran one additional model to predict the memorability scores of Exp. 1 and 2 together. This was a mixed effects linear regression model, containing all low-level features, gender, and dialect as predictors, with speaker and sentence number as random effects. This model had an adjusted *R^2^* of 0.12. Several of the low-level predictors that came out as significant overlapped with the significant features in the individual sentence models (**Supplementary Table 3**). The results of this combined model provide yet more evidence for the intrinsic memorability effect of speakers themselves across utterances, which can be reliably predicted by quantifiable acoustic features.

## Discussion

In this study, we asked, for the first time, whether intrinsic memorability exists outside the visual domain, by investigating the memorability of voices. We found that participants did in fact remember and forget the same voice clips as one another in Experiments 1 and 2, and that memory performance for the same speakers was consistent across different utterances, even within the same participants in Experiment 3. Regression models containing acoustic features were able to significantly predict the memorability of voices. Importantly, these models cross-validated out-of-sample, implying an intrinsic memorability of speakers themselves that can be reliably predicted by a combination of voice features.

These results contain both similarities and differences with previous findings in the visual memorability literature, specifically face memorability. The voice stimuli used in the current experiment exhibited a significantly consistent memorability effect across participants, as has been shown repeatedly for faces (Bainbridge et al., 2013; Bainbridge & Rissman, 2018). Memorability was also consistent across both utterances—in other words, the most memorable and forgettable speakers of the first sentence tended to be the most memorable and forgettable speakers of the second sentence. This mirrors the finding that facial identities maintain consistent memorability across different expressions and viewpoints (Bainbridge, 2017). Thus, our results provide additional evidence that intrinsic memorability exists not just for individual stimuli but for entire identities. Furthermore, these similarities between auditory and visual memorability imply the existence of a robust, multisensory property of stimuli that may also exist for taste, smell, and touch.

However, we observed differences with prior face memorability findings as well. The consistency of *d’* for voice clips reported here (average Spearman-Brown corrected ρ = 0.28) was qualitatively lower than that previously found for face images (Spearman-Brown corrected ρ = 0.82; Bainbridge et al., 2013). Thus, observers appear to be less similar to each other in their memory for voices than for faces. Furthermore, average memory performance was lower in the current study for voice clips (average *d’* = 0.31) than previously reported for face images (average *d’* = 1.11; Bainbridge et al., 2013), primarily resulting from relatively high false alarm rates for the voice stimuli. Average *d’* and split-half consistency of *d’* were also lower in the memory task for different utterances of the same speakers (Exp. 3) compared with an equivalent face memory task (Bainbridge, 2017). These results mirror previous findings suggesting that auditory long-term memory is inferior to visual long-term memory across different task designs (Bigelow & Poremba, 2014; Cohen et al., 2009; Yarmey et al., 1994). This may be explained by inherent differences between the stimulus types, in that voice clips are dynamic and contain information on speech content in addition to identity information, which may make it more difficult to recognize voices than faces. Modality-specific memory regions in the brain may also contribute to this difference; for example, it has been suggested that certain subregions of the medial temporal lobe may not support auditory memory to the same extent as visual memory (Fritz et al., 2005; Munoz-Lopez et al., 2010; Peters et al., 2007). Future studies will be needed to shed light on the role of different brain regions in memory, and sensitivity to memorability, across sensory domains.

In attempting to predict the memorability of our voice stimuli, regression models for both sentences were significantly predictive of *d’*, and even generalized out-of-sample to predict *d’* of the other sentence with above-chance accuracy. A third model was able to significantly predict memorability regardless of the specific utterance, and contained several of the same significant predictors as the first two models. Across all three models, features related to the fundamental frequency and the higher formants (e.g., F0, SD of F0, F3, SD of B3, B4, SD of F4, SD of B4) came out as significant. F0 is the perceived base pitch of a voice, and therefore conveys information about a speaker’s gender, emotion, tone, and physiology (Bishop & Keating, 2012). F0 and the third and fourth formants are thought to be important in speaker identification, as they are indicative of an individual’s voice characteristics and show high variability across speakers (Nusbaum & Magnuson, 1997; Zhang et al., 2006; Zhou et al., 2008). Also significant in multiple models were root mean square energy, which is the perceived loudness of a speech signal, and duration, which represents the tempo of speech and varies with gender, age, and dialect (Jacewicz et al., 2010). It is important to note, however, that although our models cross-validated and shared several of the same predictors, many individual predictors differed across models. Relatedly, prior research has also failed to identify a single set of acoustic features that reliably supports speaker identification, perhaps because different features may be important for identifying different speakers (Schweinberger et al., 2014; Van Lancker et al., 1985). A similar explanation may apply here, in that certain features may contribute more to the memorability of some speakers over others. Regardless, our results reveal that a combination of acoustic properties can reliably predict the memorability of voices. This finding suggests exciting avenues for the production of deep neural networks (DNNs) to predict auditory memorability, as has already been achieved with high accuracy for naturalistic images (Needell & Bainbridge, 2022).

Interestingly, the significant predictors in all three models included only low-level voice features. This contrasts with findings from the visual memorability literature, which suggest a greater influence of high-level, semantic features in determining what we remember (Kramer et al, 2023; Xie et al., 2022). Specifically, Kramer et al. found that semantic dimensions, such as animacy and whether something was a tool, explained far more variance in object memorability than visual dimensions, such as color and size. One possible explanation for this difference may come from our observed lack of consistency in the ratings of speaker personality traits. It appears that, compared with faces, participants may not agree on their judgments of subjective personality measures from hearing a voice clip (Tompkinson et al., 2023). The variability in these high-level attribute ratings may explain why they were not significantly predictive of memorability. Another explanation could be that low- and high-level features are more closely related in audition than in vision. For example, as discussed above, F0 is closely associated with the gender of a speaker, and speech tempo varies with age and dialect. Thus, it is possible that both low- and high-level features are predictive of memorability, but due to their multicollinearity, only low-level predictors remained in our models. However, it is important to note that regardless of this multicollinearity, low-level features were consistently selected by the models over the related high-level features.

Our models of voice memorability could be useful for researchers to control for the memorability of their stimuli in auditory experiments. In daily life, these findings could be used by teachers to select memorable educational videos, by language-learning apps to boost the memorability of their virtual instructors, and by nonprofits to produce memorable advertisements highlighting important causes. Particularly memorable virtual assistants could help individuals plan their schedules and carry out daily tasks, which may be especially beneficial for patients with Alzheimer’s Disease or other memory deficits. Overall, our results prove that we are more alike than we may think in the voices we remember.

## Methods

### Participants

Participants were recruited through the online platform Amazon Mechanical Turk (AMT). To participate in the study, participants had to have previously completed ≥ 50 AMT Human Intelligence Tasks (“HIT”s), have an AMT approval rating of ≥ 95%, and reside in the U.S. 3930 participants performed Experiment 1, 4373 participants performed Experiment 2, and 3761 participants performed Experiment 3. Following our exclusion criteria (see below), data from 1325 participants was analyzed from Experiment 1, data from 1425 participants was analyzed from Experiment 2, and data from 1007 participants was analyzed from Experiment 3. We arrived at these sample sizes so that each target stimulus was heard by an average of at least 40 participants, as previous work finds that a stable measure of memorability is reached at 40 participant ratings/ stimulus (Isola et al., 2014). A small proportion of participants completed more than one memory experiment (N = 417), as each experiment was run three months apart. In total, data from 3340 participants was analyzed (1561 female, 1461 male, 318 other/did not answer).

In addition, another 1147 participants performed the attribute rating experiment. Following our exclusion criteria, data from 903 participants was analyzed (469 female, 426 male, 8 other/did not answer). We arrived at this sample size to collect 30 ratings for each attribute antonym-pair, following the procedure in Bainbridge et al., 2013. Compensation was provided at a rate of $6 per hour in all experiments. Participants gave electronic informed consent prior to the experiment, following the University of Chicago Institutional Review Board guidelines (IRB19-1395).

### Stimuli

Audio stimuli came from the DARPA TIMIT Acoustic-Phonetic Continuous Speech Corpus (“TIMIT”), a large-scale database containing recordings of 630 speakers (483 male, 192 female) from eight major U.S. dialects, speaking phonetically-diverse sentences (Garofolo et al., 1993). The represented dialects include: New England, Northern, North Midland, South Midland, Southern, New York City, Western, and Army Brat. The TIMIT corpus contains the same two sentences spoken by all 630 speakers. The first sentence (“*She had your dark suit in greasy wash water all year*”) was used as the stimuli in Experiment 1, the second sentence (“*Don’t ask me to carry an oily rag like that*”) was used as the stimuli in Experiment 2, and both sentences were used as the stimuli in Experiment 3. Because of the imbalance in gender of the speakers in the database, we subsampled from the set of male speakers to equal the number of female speakers. We also identified voice clips greater than five seconds in length, and excluded both sentences spoken by these speakers (three total speakers). This resulted in clips of 378 speakers speaking both Sentence 1 and 2 (189 male, 189 female; 34 New England, 60 Northern, 55 North Midland, 56 South Midland, 68 Southern, 31 New York City, 51 Western, 23 Army Brat dialect). The average length of Sentence 1 was 3.42 seconds (min = 2.35, max = 4.99, SD = 0.45), and the average length of Sentence 2 was 2.81 seconds (min = 2.01, max = 4.46, SD = 0.41).

### Memory experiments

Experiment 3 of this study was preregistered on the Open Science Framework (https://osf.io/pm8wa). All three memory experiments followed the same general procedure. Prior to beginning the main experiment, participants answered two common sense questions. Those who did not correctly answer both questions were replaced with new participants. Participants also rated the noise level of their surroundings on a scale from 1-5. Those who selected 4 or 5 were replaced with new participants. Lastly, participants transcribed a practice sentence that differed from the sentence in the main task and was randomized across participants. Those who did not include at least two correct words in their answer were replaced with new participants. These checks were included to ensure that participants were paying attention, and were able to listen to audio stimuli for the main task.

The main experiment was a continuous recognition task in which participants listened to a randomized sequence of speakers. Experiment 1 contained the first sentence as stimuli, Experiment 2 contained the second sentence as stimuli, and Experiment 3 contained both sentences. Voice clips were played within a five-second-long stimulus interval, each separated by two seconds of pink (1/*f*) noise (**Figure 1**). Pink noise was chosen for the interstimulus interval as it is representative of the spectrum of natural sounds (Szendro et al., 2001). In Experiments 1 and 2, ∼56 stimuli were randomly selected to make up each participant’s stimulus sequence. ∼17 of these stimuli were randomly selected as target stimuli, and the remaining ∼39 were used as filler stimuli. (Due to the randomized generation of the stimulus sequence, these values could vary slightly across participants.) All stimuli served as targets for some participants and fillers for others. Target stimuli could repeat once 16 to 24 trials after the first presentation (on average, two minutes and 20 seconds later). Filler stimuli could repeat much more frequently, one to nine trials after the first presentation (on average, 35 seconds later). Participants were told to press the ‘r’ key on their keyboard whenever they heard a repeated voice clip. In Experiment 3, repeats were of the same speaker, speaking the other sentence. For example, if a repeated speaker was first played in the sequence speaking Sentence 1, they would later be played speaking Sentence 2, and vice versa. Here, participants were told to press the ‘r’ key whenever they heard a repeated *speaker*, even across different sentences. To account for the increased difficulty of the task, the number of unique stimuli and the timing between repeats in Experiment 3 was slightly updated. ∼47 stimuli were randomly selected to make up each participant’s sequence, with ∼22 being target stimuli and ∼25 being filler stimuli. Target repeats occurred nine to 15 trials after the first presentation (on average, 84 seconds later), and filler repeats occurred one to seven trials after the first presentation (on average, 28 seconds later). Although participants responded to both target and filler repeats, the fillers were analyzed only as a vigilance check—participants who missed more than 70% of filler repeats or had false alarms on more than 70% of the initial presentation of fillers were excluded from further analysis. In total, the task took approximately 10 minutes to complete.

### Attribute rating experiment

A separate group of AMT participants were recruited to perform the attribute rating experiment. Here, participants listened to the same Sentence 1 stimuli used in Experiment 1, and rated them according to 14 personality-related attributes and their antonyms. These attributes included *attractive/unattractive, calm/aggressive, competent/incompetent, confident/uncertain, dominant/submissive, expressive/unemotional, familiar/unfamiliar, likable/unlikable, masculine/feminine, memorable/forgettable, trustworthy/untrustworthy, typical/atypical, warm/cold,* and *young/old*. These traits were borrowed from previous work on memorability and personality judgments of faces (Bainbridge et al., 2013; Oosterhof & Todorov, 2008; Vokey & Read, 1992), and personality judgments of voices (McAleer et al., 2014). As these attributes did not end up being significantly predictive of memorability in our regression model, we did not collect ratings for the Sentence 2 stimuli.

In this task, participants were presented with a single voice clip, which they could replay multiple times. They were asked to rate the clip according to how much the speaker’s voice sounded like each of the attributes listed above (e.g., “How *likable* does this voice sound?”). We ran two versions of this survey on separate groups of participants, with the attributes and their antonyms randomly split across versions. Participants rated the voice clip according to each of the 14 attributes on a 9-point Likert scale, from 1 (“Not at all”) to 9 (“Extremely”). If they needed to, they could click a pop-up window next to each attribute to read a dictionary definition of the word. In addition, we included a catch question in which participants selected the “*n*th” word in the spoken sentence, with the word number randomized across participants. Those who incorrectly answered this question were replaced with new participants. Participants were able to take the survey multiple times (on average, 13 times per participant), each time rating a different voice clip according to the same 14 attributes. In total, 15 ratings of each attribute were collected for each voice clip.

## Analyses

### Consistency of stimulus memorability across participants

We calculated the hit rate (HR) of each speaker as the proportion of target repeats that were correctly identified by participants. We also calculated the false alarm rate (FAR) of each speaker as the proportion of the initial presentation of targets that participants incorrectly identified as a repeat. From these two measures, we calculated the *d’* of each speaker as *norminv(HR) - norminv(FAR)*, which served as our main measure of memorability. These measures were calculatedly separately for all three memory experiments.

To measure the consistency across participants in the speakers they (correctly and incorrectly) remembered and forgot, we ran a split-half consistency analysis on *d’* values (Bainbridge et al., 2013; Isola et al., 2011), again for all three experiments separately. In this analysis, the participant pool is randomly split into two equal groups, and average *d’* is computed for all stimuli in each group. The Spearman rank correlation is then measured between *d’* values of group 1 and group 2. Chance level correlation is determined by correlating group 1 with a shuffled group 2. This is repeated over 1000 iterations, after which the resulting Spearman rho values are averaged to produce the final correlation. The significance of this value is determined using a one-tailed nonparametric permutation test, comparing the average correlation value with the 1000 shuffled correlation values. A significant correlation indicates that participants are consistent in their memory performance for stimuli. To account for the fact that this analysis reduces the sample size by half, we performed the Spearman-Brown split-half reliability correction on the average correlation value, computed as (2 * ρ) / (1 + ρ).

### Low-level features analysis

We aimed to predict the memorability scores of the stimuli in Experiments 1 and 2 from a combination of both low-level and high-level voice features. To measure low-level features, we used VoiceSauce, an application implemented in MATLAB that computes 44 different acoustic features of a given voice recording (Shue et al., 2011). VoiceSauce calculates several of these features multiple times according to different algorithms, including the Straight algorithm (prefix abbreviated “str”; Kawahara et al., 1998), the Snack Sound Toolkit (“s”; Sjölander, 2004), Praat (“p”; Boersma & Weenink, 2023) and Sun’s Subharmonic-to-Harmonic Ratio method (“shr”; Sun, 2002). The full list of features calculated by VoiceSauce includes the fundamental frequency (strF0, sF0, pF0, shrF0), frequency of the first four formants (sF1, sF2, sF3, sF4, pF1, pF2, pF3, pF4), frequency of the harmonic nearest 2000 Hz (F2K), bandwidth of the first four formants (sB1, sB2, sB3, sB4, pB1, pB2, pB3, pB4), amplitude of the first, second, and fourth harmonics (H1, H2, H4), amplitude of the harmonics nearest the first three formants (A1, A2, A3), amplitude of the harmonic nearest 2000 Hz (H2K), amplitude of the harmonic nearest 5000 Hz (H5K), amplitude difference between the first and second harmonic (H1-H2), amplitude difference between the second and fourth harmonic (H2-H4), amplitude difference between the first harmonic and the harmonic nearest the first formant (H1-A1), amplitude difference between the first harmonic and the harmonic nearest the second formant (H1-A2), amplitude difference between the first harmonic and the harmonic nearest the third formant (H1-A3), amplitude difference between the fourth harmonic and the harmonic nearest 2000 Hz (H4-H2K), amplitude difference between the harmonic nearest 2000 Hz and the harmonic nearest 5000 Hz (H2K-H5K), root mean square energy, cepstral peak prominence (CPP), harmonic-to-noise ratio between 0 and 500 Hz (HNR05), harmonic-to-noise ratio between 0 and 1500 Hz (HNR15), harmonic-to-noise ratio between 0 and 2500 Hz (HNR25), harmonic-to-noise ratio between 0 and 3500 Hz (HNR35), subharmonic-to-harmonic ratio (SHR), and strength of excitation (SoE). Because VoiceSauce computes these measures for every millisecond of a voice clip, we computed one average measure of each feature for all voice clips of both sentences. We also computed the standard deviation of each feature across the voice clip, as a measure of how much the feature varied over the sentence. Thus, we considered 88 low-level acoustic features in total (44 mean measures and 44 standard deviation measures). We also included the duration of the voice clips as a low-level feature. Duration can be thought of as inversely related to the speed of a speaker’s speech, because all clips were composed of the same sentence.

### High-level features analysis

Using the results of the attribute rating experiment, we first correlated participants’ ratings of the attribute antonym pairs using the Pearson correlation. This step tested for across-participant consistency in ratings, as ratings of antonyms were expected to be negatively correlated with each other. We then aligned the ratings of the antonym pairs onto the same scale by subtracting ratings of corresponding antonyms from 10. For example, a voice originally rated as 1 on “trustworthiness” would be rated as 9 on “untrustworthiness.” This resulted in 30 ratings per combined attribute. We ran a split-half consistency analysis on these ratings for each attribute, as described above, to test for consistency in participants’ subjective judgments of voices. Finally, we averaged across the 30 ratings for each of the 14 attributes for all Sentence 1 voice clips.

### Linear regression models of memorability

In preparing to run our models predicting voice memorability, we normalized all continuous variables into standardized z-scores. For the Sentence 1 stimuli, this included the mean and standard deviation measures of the 44 low-level features, the “duration” measure, and the 14 high-level attributes, while for the Sentence 2 stimuli, this included only the mean and standard deviation of the low-level features, as well as duration. In addition to these variables, we also included two categorical variables: the gender and dialect of the speakers.

We used stepwise linear regression (*stepwiselm* in MATLAB) to arrive at a multiple regression model predicting voice *d’*, separately for the Sentence 1 and 2 stimuli from Experiment 1 and 2, respectively. This function starts with a constant model and adds or removes terms one-by-one to optimize the specified criterion. The selected criterion was “SSE”, or the *p*-value for an *F*-test of the change in the sum of squared error that results from adding or removing a given term. The final model arrived at by the regression function is therefore the most significantly predictive model, from all possible combinations of predictors. For both the Sentence 1 and Sentence 2 final models, we ran an *F*-test to test the overall significance of the model, and used adjusted R^2^ as our measure of model fit.

We aimed to see whether these models cross-validated to the other sentence by the same speakers. To do so, we ran the Sentence 1 model formula on the measures of the same features of the Sentence 2 voice clips to predict the memorability of the Sentence 2 stimuli, and vice versa, using the Sentence 2 model to predict Sentence 1 memorability. We then took the Spearman correlation between those predicted scores and the true *d’* scores for both Sentence 1 and Sentence 2 stimuli, and tested for significance of the correlation value.

Finally, we ran one additional model to predict the *d’* of all voice clips across both experiments. This was a mixed effects linear regression model (*fitlme* in MATLAB), which included all normalized low-level features, gender, and dialect as predictors. Intercepts for speaker and sentence number were included as random effects. Adjusted R^2^ was again used as the measure of model fit.

## Acknowledgements

We would like to thank Kristin Van Engen, Howard Nusbaum, and Marc Berman for their insightful feedback on the analyses and the work as a whole. This research was supported by the National Science Foundation under Grant No. 2329776 awarded to W.A.B., the NSF Graduate Research Fellowship awarded to C.R., and the University of Chicago Quad Undergraduate Research Scholarship awarded to E.G. Experiment 3 of this study was preregistered on the Open Science Framework (https://osf.io/pm8wa). All data will be made publicly available upon publication at https://osf.io/pybwd/.

## Notes

### Competing Interest Statement

The authors have declared no competing interest.

