## Supplementary Material for "The Memorability of Voices is Predictable and Consistent across Listeners"

Correspondence to: Cambria Revsine

301 Green Hall

5848 S. University Ave.

Chicago, IL 60637

Figure/Table Page

**Supplementary Figure 1:** Correlation matrix of Sentence 2 low-level features. 3

**Supplementary Table 1:** Stepwise regression model predicting Sentence 1 *d’* from Experiment 1. 4

**Supplementary Table 2:** Stepwise regression model predicting Sentence 2 *d’* from Experiment 2. 5

**Supplementary Table 3:** Mixed effects regression model predicting *d’* of both sentences from Experiment 1 and 2. 6

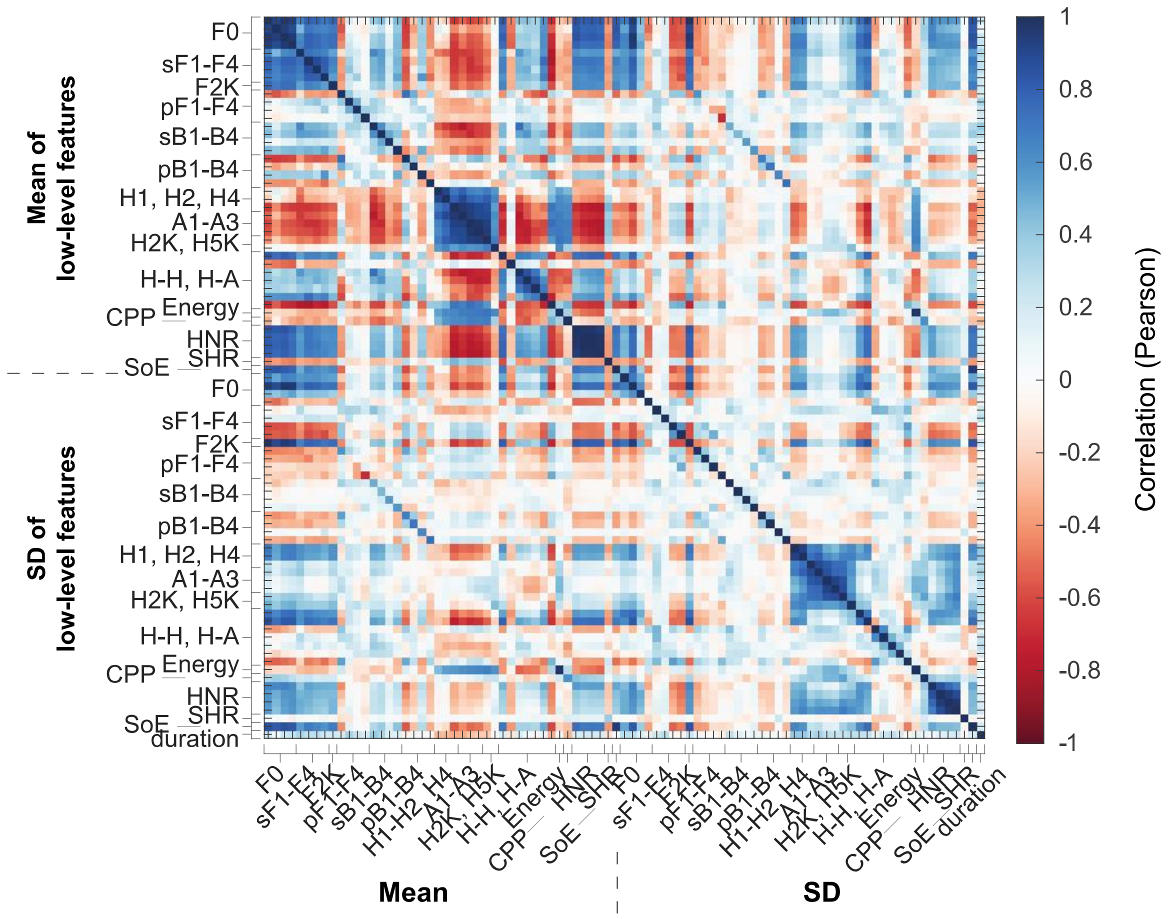


**Supplementary Figure 1.** Correlation matrix of Sentence 2 low-level features. Similar to Figure 4, this matrix contains mean measures of the 44 low-level acoustic features of Sentence 2 stimuli, followed by the standard deviation (SD) of the same features, followed by the "duration" measure. Many strong correlations can be seen between related features.

*Sentence 1 Model*

| | **$\beta$ estimate** | **SE** | ***t*-statistic** | ***p*-value** |
|---|---|---|---|---|
| **Duration** | 0.19 | 0.05 | 3.78 | $1.9 \times 10^{-4}$ |
| **SD of pB3** | -0.15 | 0.05 | -3.06 | 0.002 |
| **Duration: SD of pB3** | -0.17 | 0.05 | -3.23 | 0.001 |
| *Model Stats:* $F(7,370) = 13.8$, $p < 0.001$; Adj. $R^2 = 0.09$ | | | | |

**Supplementary Table 1.** Stepwise regression model predicting Sentence 1 *d'* from Experiment 1. Significant model predictors include duration of the voice clip, standard deviation of the bandwidth of the 3rd formant (B3), and their interaction. A closer look at the interaction term revealed that duration had a greater effect on memorability the lower the standard deviation of B3. The beta estimate, standard error of the estimate, *t*-statistic, and *p*-value of all model predictors are reported. Model summary statistics, including results of an *F*-test and the adjusted $R^2$ value, are also reported.

*Sentence 2 Model*

| | ***β* estimate** | **SE** | ***t*-statistic** | ***p*-value** |
|---|---|---|---|---|
| **shrF0** | 0.25 | 0.06 | 4.35 | $1.8 \times 10^{-5}$ |
| **pF3** | 0.19 | 0.05 | 3.80 | $1.7 \times 10^{-4}$ |
| **SD of sF4** | 0.21 | 0.06 | 3.58 | $3.8 \times 10^{-4}$ |
| **SD of H4-H2K** | -0.15 | 0.05 | -2.90 | 0.004 |
| **Energy** | 0.26 | 0.05 | 5.29 | $2.1 \times 10^{-7}$ |
| **shrF0: pF3** | -0.14 | 0.05 | -2.68 | 0.008 |
| **shrF0: SD of H4-H2K** | 0.12 | 0.05 | 2.43 | 0.02 |
| *Model Stats:* $F(7,370) = 10.1$, $p < 0.001$ Adj. $R^2 = 0.15$ | | | | |

**Supplementary Table 2:** Stepwise regression model predicting Sentence 2 *d'* from Experiment 2. Layout of regression table as in Supplementary Table 1. Significant model predictors include the fundamental frequency (F0), frequency of the third formant (F3), standard deviation of the frequency of the fourth formant (F4), standard deviation of the amplitude difference between the $4^{th}$ harmonic and the harmonic nearest 2000 Hz (H4-H2K), and root mean square energy. The model also includes an interaction term between F0 and F3, in which F3 had a greater effect on memorability the lower the F0, and a second interaction term between F0 and SD of H4-H2K, in which SD of H4-H2K had a greater effect on memorability the higher the F0.

*Combined Sentence Model*

| | ***β* estimate** | **SE** | ***t*-statistic** | ***p*-value** |
|---|---|---|---|---|
| **strF0** | 1.44 | 0.58 | 2.48 | 0.01 |
| sF0 | 0.05 | 0.29 | 0.18 | 0.86 |
| pF0 | -0.48 | 0.30 | -1.60 | 0.11 |
| shrF0 | 0.09 | 0.26 | 0.35 | 0.73 |
| sF1 | -0.18 | 0.76 | -0.23 | 0.82 |
| sF2 | -0.51 | 1.03 | -0.49 | 0.62 |
| sF3 | -0.40 | 0.61 | -0.66 | 0.51 |
| sF4 | 0.16 | 0.12 | 1.37 | 0.17 |
| F2K | -0.03 | 0.07 | -0.39 | 0.70 |
| pF1 | -0.03 | 0.13 | -0.25 | 0.80 |
| pF2 | 0.02 | 0.10 | 0.26 | 0.80 |
| pF3 | 0.08 | 0.09 | 0.94 | 0.35 |
| pF4 | -0.05 | 0.08 | -0.59 | 0.55 |
| sB1 | 0.03 | 0.13 | 0.27 | 0.78 |
| sB2 | 0.09 | 0.13 | 0.70 | 0.48 |
| sB3 | -0.02 | 0.09 | -0.20 | 0.84 |
| **sB4** | 0.16 | 0.07 | 2.30 | 0.02 |
| pB1 | -0.01 | 0.08 | -0.16 | 0.88 |
| pB2 | 0.06 | 0.07 | 0.86 | 0.39 |
| pB3 | -0.04 | 0.07 | -0.52 | 0.60 |
| pB4 | 0.08 | 0.08 | 0.94 | 0.35 |
| **H1** | -3.45 | 1.47 | -2.34 | 0.02 |
| H2 | -0.63 | 0.39 | -1.62 | 0.11 |
| **H4** | -0.93 | 0.33 | -2.85 | 0.004 |
| A1 | 4.67 | 2.57 | 1.81 | 0.07 |
| A2 | -2.89 | 3.41 | -0.85 | 0.40 |
| A3 | -0.50 | 0.87 | -0.58 | 0.56 |
| H2K | 0.34 | 0.25 | 1.39 | 0.17 |

| | | | | |
|---|---|---|---|---|
| H5K | 3.59 | 2.20 | 1.63 | 0.10 |
| H1-H2 | 1.90 | 1.30 | 1.46 | 0.14 |
| H2-H4 | 1.69 | 1.02 | 1.66 | 0.10 |
| H1-A1 | 2.30 | 1.32 | 1.75 | 0.08 |
| H1-A2 | -1.75 | 2.12 | -0.82 | 0.41 |
| H1-A3 | -0.56 | 0.59 | -0.96 | 0.34 |
| H4-H2K | 3.10 | 1.80 | 1.72 | 0.09 |
| H2K-H5K | 5.22 | 3.12 | 1.67 | 0.09 |
| **Energy** | 0.44 | 0.19 | 2.28 | 0.02 |
| CPP | -0.27 | 0.14 | -1.92 | 0.05 |
| HNR05 | -0.09 | 0.57 | -0.16 | 0.87 |
| HNR15 | -0.48 | 0.95 | -0.50 | 0.61 |
| HNR25 | 0.59 | 1.25 | 0.47 | 0.64 |
| HNR35 | 0.41 | 0.77 | 0.53 | 0.60 |
| SHR | 0.07 | 0.06 | 1.20 | 0.23 |
| SoE | 0.02 | 0.19 | 0.09 | 0.93 |
| SD of strF0 | 0.07 | 0.07 | 1.01 | 0.31 |
| **SD of sF0** | 0.52 | 0.22 | 2.40 | 0.02 |
| **SD of pF0** | -0.10 | 0.05 | -2.01 | 0.04 |
| SD of shrF0 | -0.04 | 0.07 | -0.55 | 0.58 |
| SD of sF1 | 0.06 | 0.10 | 0.63 | 0.53 |
| SD of sF2 | -0.02 | 0.09 | -0.21 | 0.84 |
| SD of sF3 | -0.02 | 0.08 | -0.23 | 0.82 |
| **SD of sF4** | 0.14 | 0.06 | 2.36 | 0.02 |
| SD of F2K | -0.35 | 0.30 | -1.17 | 0.24 |
| SD of pF1 | 0.03 | 0.12 | 0.25 | 0.80 |
| SD of pF2 | -0.11 | 0.08 | -1.39 | 0.17 |
| SD of pF3 | -0.05 | 0.06 | -0.90 | 0.37 |
| SD of pF4 | -0.06 | 0.08 | -0.81 | 0.42 |
| SD of sB1 | 0.01 | 0.07 | 0.13 | 0.89 |
| **SD of sB2** | -0.15 | 0.06 | -2.79 | 0.005 |

| | | | | |
|---|---|---|---|---|
| SD of sB3 | 0.05 | 0.05 | 1.03 | 0.30 |
| SD of sB4 | -0.05 | 0.05 | -0.94 | 0.35 |
| SD of pB1 | 0.08 | 0.06 | 1.32 | 0.19 |
| SD of pB2 | -0.08 | 0.07 | -1.26 | 0.21 |
| SD of pB3 | -0.07 | 0.05 | -1.37 | 0.17 |
| SD of pB4 | -0.04 | 0.06 | -0.70 | 0.48 |
| SD of H1 | 0.06 | 0.15 | 0.37 | 0.71 |
| SD of H2 | -0.13 | 0.16 | -0.77 | 0.44 |
| SD of H4 | 0.24 | 0.13 | 1.77 | 0.08 |
| SD of A1 | -0.35 | 0.19 | -1.86 | 0.06 |
| SD of A2 | 0.26 | 0.17 | 1.56 | 0.12 |
| SD of A3 | -0.05 | 0.13 | -0.34 | 0.73 |
| SD of H2K | -0.13 | 0.13 | -1.02 | 0.31 |
| **SD of H5K** | 0.14 | 0.06 | 2.14 | 0.03 |
| SD of H1-H2 | -0.07 | 0.08 | -0.86 | 0.39 |
| SD of H2-H4 | -0.07 | 0.10 | -0.74 | 0.46 |
| SD of H1-A1 | 0.14 | 0.10 | 1.41 | 0.16 |
| SD of H1-A2 | 0.02 | 0.07 | 0.31 | 0.76 |
| SD of H1-A3 | -0.03 | 0.07 | -0.42 | 0.67 |
| SD of H4-H2K | -0.08 | 0.05 | -1.54 | 0.12 |
| SD of H2K-H5K | -0.06 | 0.06 | -0.85 | 0.40 |
| SD of Energy | -0.21 | 0.17 | -1.22 | 0.22 |
| SD of CPP | 0.03 | 0.10 | 0.30 | 0.77 |
| **SD of HNR05** | 0.20 | 0.10 | 2.06 | 0.04 |
| **SD of HNR15** | -0.51 | 0.17 | -3.03 | 0.003 |
| SD of HNR25 | 0.55 | 0.31 | 1.78 | 0.08 |
| SD of HNR35 | -0.22 | 0.22 | -1.03 | 0.31 |
| SD of SHR | -0.06 | 0.05 | -1.24 | 0.21 |
| SD of SoE | -0.08 | 0.21 | -0.37 | 0.71 |
| **Duration** | 0.16 | 0.06 | 2.79 | 0.006 |
| Female | -0.12 | 0.24 | -0.50 | 0.62 |

| | | | | |
|---|---|---|---|---|
| Northern | -0.08 | 0.15 | -0.51 | 0.61 |
| North Midland | 0.14 | 0.15 | 0.92 | 0.36 |
| South Midland | 0.24 | 0.15 | 1.58 | 0.11 |
| Southern | 0.17 | 0.15 | 1.17 | 0.24 |
| New York City | -0.01 | 0.17 | -0.03 | 0.98 |
| Western | 0.13 | 0.16 | 0.86 | 0.39 |
| Army Brat | -0.08 | 0.19 | -0.43 | 0.67 |
| *Model Stats:* Adj. $R^2 = 0.12$ | | | | |

**Supplementary Table 3:** Mixed effects regression model predicting *d'* of both sentences from Experiment 1 and 2. Layout of regression table as in Supplementary Table 1 and 2. Unlike the previous fixed effects models resulting from stepwise regression, this model includes all low-level features, gender, and dialect as predictors, as well as speaker and sentence number as random intercepts. Significant predictors, in bold, include the fundamental frequency (F0), bandwidth of the fourth formant (B4), amplitude of the first harmonic (H1), amplitude of the fourth harmonic (H4), root mean square energy, standard deviation of F0, standard deviation of F4, standard deviation of B2, standard deviation of the amplitude of the harmonic nearest 5000 Hz (H5K), standard deviation of the harmonic-to-noise ratio between 0 and 500 Hz (HNR05), standard deviation of the harmonic-to-noise ratio between 0 and 1500 Hz (HNR15), and duration of the voice clip. Several of these significant predictors, including F0, energy, SD of F4, and duration, also appear in the individual sentence models.
